# Structural Model for Self-Limiting β-strand Arrangement Within an Alzheimer’s Amyloid-β Oligomer

**DOI:** 10.1101/2022.12.06.519347

**Authors:** Yuan Gao, Ramesh Prasad, Peter S. Randolph, Jens O. Watzlawik, Alicia S. Robang, Cong Guo, Scott M. Stagg, Huan-Xiang Zhou, Terrone L. Rosenberry, Anant K. Paravastu

**Affiliations:** School of Chemical and Biomolecular Engineering, Georgia Institute of Technology, 311 Ferst Drive NW, Atlanta, GA 30332, USA; Department of Chemistry, University of Illinois at Chicago, Chicago, IL 60607, USA; Department of Physics, University of Illinois at Chicago, Chicago, IL 60607, USA; Institute of Molecular Biophysics, Florida State University, Tallahassee, FL 32306, USA; Department of Biological Sciences, Florida State University, Tallahassee, FL 32306, USA; Departments of Neuroscience and Pharmacology, Mayo Clinic College of Medicine, 4500 San Pablo Road, Jacksonville, FL 32224, USA; Department of Physics and International Centre for Quantum and Molecular Structures, Shanghai University, 99 Shangda Road, Shanghai, China

## Abstract

Previous reports revealed that sodium dodecyl sulfate near its critical micelle concentration can drive the assembly of Aβ42 along an oligomeric pathway. This pathway produces a 150 kDa peptide oligomer (approximately 32 peptide molecules or protomers) that does not aggregate further into amyloid fibrils. Solid-state nuclear magnetic resonance (NMR) spectroscopy revealed structural features distinguishing the 150 kDa oligomer from fibrils. A puzzling feature was the coexistence of parallel and antiparallel β-sheets within the oligomer structure. Here we present new atomic-level structural constraints obtained via solid-state NMR spectroscopy, benefitting from improved resolution via sample concentration by ultracentrifugation. In addition, two-dimensional cryo-electron microscopy (cryo-EM) reconstruction revealed a 4-fold symmetric shape. We propose a structural model to rationalize the solid-sate NMR- and cryo-EM-derived structural constraints. This model has a hollow square cylinder shape, with antiparallel β-sheets formed by residues 33-39 lining the inner walls and parallel β-sheets formed by residues 11-22 lining the outer walls. Within successive layers, the outer β-strands on each side of the square cylinder alternate between two forms: one within a U-shaped protomer and another within L-shaped protomer. Molecular dynamics simulations show that, when the oligomer model is embedded in a lipid membrane, ions permeate through the central pore, with cation selectivity. The model further motivates an assembly pathway-based interpretation that may explain why the 150 kDa oligomer does not undergo further aggregation into amyloid fibrils.

**Significance Statement:** Aβ oligomers are thought to be the most toxic species in Alzheimer’s disease. Their sizes range from 2 to ∼50 protomers. Most published experimental data on Aβ oligomers indicate that they, like fibrils, are composed of β-sheets, but it is a mystery why any β-sheet aggregate would exist as a stable oligomer without undergoing further aggregation into fibrils. Here, structural constraints from solid-state NMR and cryo-EM led us to an oligomer model with a hollow square cylinder shape capable of conducting ions when embedded in a lipid membrane. Based on the model, we argue that geometric frustration may distinguish the assembly pathway that produces this oligomer from fibril-forming assembly pathways.

## Introduction

An unmet challenge in Alzheimer’s disease research is the determination of structures of amyloid-β peptide (Aβ) oligomers. Oligomers are the smallest peptide aggregates, composed of 2 to ∼50 Aβ protomers, and are considered the most toxic species in patient brains (1-5). The small size of oligomers may matter as it, e.g., may correspond to higher diffusivities and more interactions with neuronal membranes (6). Understanding oligomeric structures, their underlying assembly mechanisms, and oligomer-membrane interactions could help elucidate possible mechanisms of oligomer toxicity. Proposed mechanisms include outer and mitochondrial membrane permeabilization (7-9), disruption of Na^+^ or Ca^2+^ regulation (10, 11), and receptor-mediated apoptosis (12, 13). Furthermore, oligomers could assemble in the neuronal membrane following enzymatic cleavage of the amyloid precursor protein, where they could become pathological without being readily detectable (14, 15).

The structural biology of oligomers is challenging because it is difficult to produce oligomeric samples with homogeneous and stable structures. Nevertheless, studies of Aβ aggregation have revealed relevant insight: Aβ can undergo multiple assembly pathways to create various possible structures (16-23), and these pathways are susceptible to Aβ mutations (19), the solution environment (24), and interactions with interfaces (25). Environment-dependent pathways are likely to affect structural distributions for oligomeric and protofibrillar assemblies. While there are methods to purify specific fibril structures (e.g., seeded propagation) (16, 26), such approaches may not apply to protofibrils or oligomers. Despite challenges, some reports describe oligomer structural characterization, primarily for sizes ranging from dimers to hexamers (27-32).

Our previous work established critical information on a 150 kDa oligomer (∼32 Aβ protomers) formed from initial exposure to sodium dodecyl sulfate (SDS) micelles (33). The 150 kDa oligomers are formed by the 42-residue isoform of Aβ (Aβ42) and not by the 40-resdiue isoform (Aβ40). We demonstrated repeatable experimental structural data on the 150 kDa oligomer and mysterious structural constraints (34, 35). Presently unexplained structural constraints include the coexistence of parallel β-sheets formed by residues E11-V24 (the “N-strand”) and antiparallel β-sheets formed by residues A30-A42 (the “C-strand”) and a complex pattern of proximities of N-strands and C-strand residues (34). Supplementary Figure S1 shows that the pattern of inter-residue proximities observed for the 150 kDa oligomer is inconsistent with previously published structures and structural models for Aβ aggregates. It is also unknown why the 150 kDa oligomer is “off-pathway” to fibril formation (33, 36): this oligomer does not undergo further aggregation to form fibrils or accelerate the assembly of fibrils when added to initially monomeric Aβ42 solutions (37, 38).

In this contribution, we construct a structural model for the 150 kDa oligomer to rationalize known structural features and new constraints. New constraints include those derived from solid-state nuclear magnetic resonance (NMR) spectroscopy and cryo-electron microscopy (cryo-EM). To obtain new structural information, we show that NMR spectral resolution can be improved by avoiding lyophilization using ultracentrifugation to pellet oligomers into solid-state NMR rotors (sample holders). Cryo-EM measurements provide data on oligomer size and shape via 2D reconstruction. Based on our structural model, we propose that oligomer size could be limited by geometric frustration in the β-strand organization.

## Results

### Constraints on molecular conformation, β-strand alignment, and multisite occupancy from solid-state NMR

Figure 1A presents observed (symbols) and predicted (colors) inter-residue proximities in a “contact chart.” The symbols defined in the next paragraph indicate which pairs of residues were revealed by 2D NMR to be spatially proximate in the 150 kDa oligomer; each row and column in the chart corresponds to a specific residue within Aβ42. The NMR experiments employed selective ^13^C labeling for unequivocal crosspeak assignment; the labeling schemes for 17 samples analyzed are listed in Supplementary Table S1. The NMR data are shown in in Figures 1B, Supplementary Figure S2, and Supplementary Table S2. Inter-residue contacts were detected via inter-residue off-diagonal peaks (crosspeaks) in 2D ^13^C-^13^C NMR spectra. The near-diagonal region of the contact chart is shaded gray because contacts between residues *i* and *i* ± 2 near-diagonal contacts are expected for β-strands regardless of inter-strand arrangement. The color scale corresponds to two incomplete model β-sheets known to occur within this structure (see Supplementary Figure S3). As we showed previously (35), residues 10-24 form the “N-strand,” which arranges into an out-of-register parallel β-sheet. Residues 30-42 form the “C-strand,” which arranges into an antiparallel β-sheet (34). Since each of these models only considers one β-strand region within the oligomer, they are incapable of rationalizing the pattern of contacts between N-strand residues and C-strand residues or contacts involving residues G25-G29. The goal of the present contribution is to explain the entirety of Figure 1A with a structural model.

**Figure 1:**
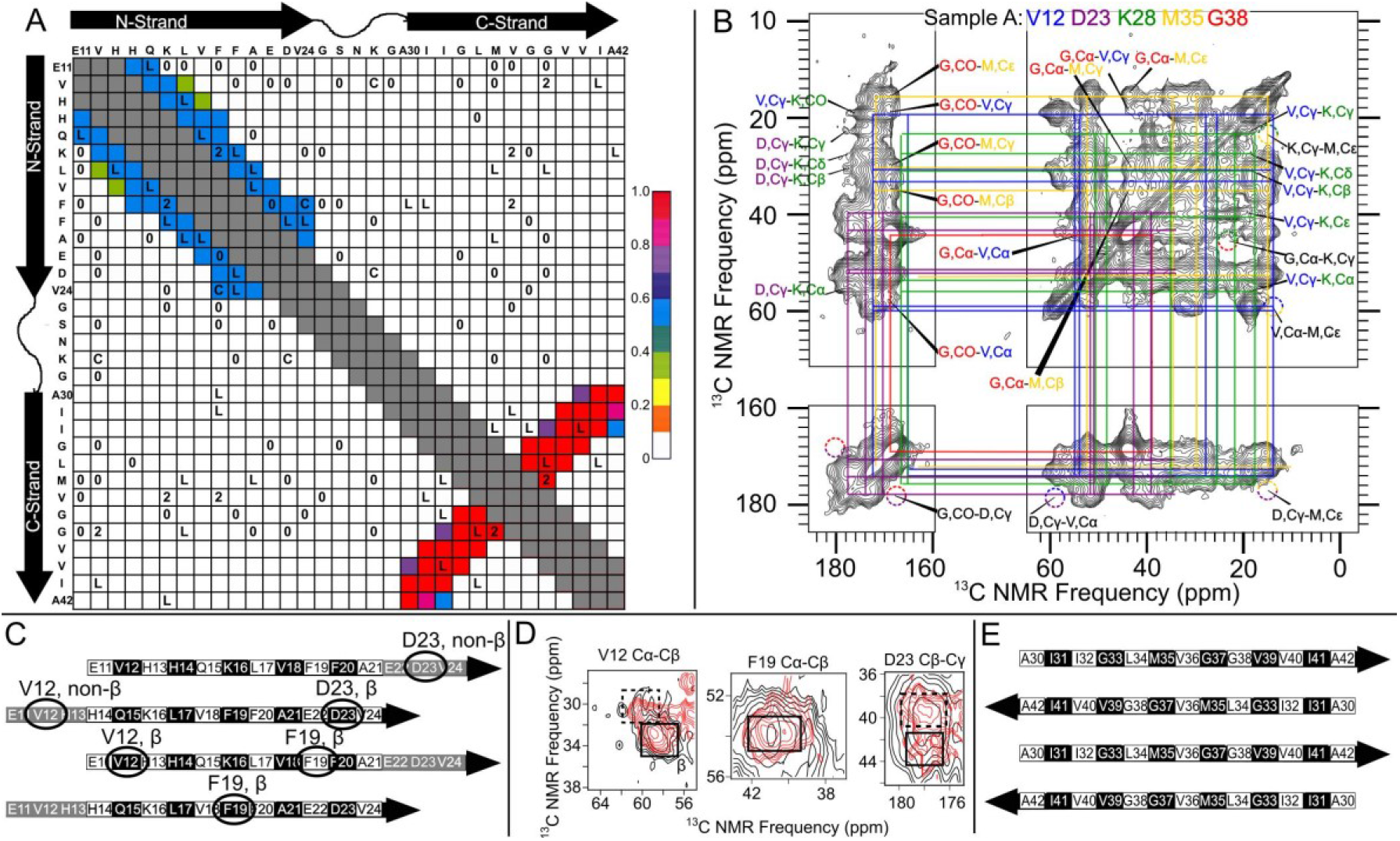
A) Contact chart describing the pattern of spatially proximate residues detected experimentally by 2D NMR (symbols) and predicted by molecular structural models (colors). The L, C, and 2 symbols indicate inter-residue contacts detected using lyophilized samples, centrifuged samples, or both, respectively. The color scale indicates the fraction of nearest-neighbor combinations of residues predicted to contribute to detectable 2D NMR contacts, using the single-β-sheet models in Figure S3. B) A 2D DARR ^13^C-^13^C NMR spectrum from Sample A. The colored lines indicate spectral assignments via intra-residue crosspeaks. Arrows and text mark observed inter-residue crosspeaks. Dashed circles show regions where crosspeaks might be expected but were not detected, indicating residue combinations that yielded “0” symbols in panel A. C) Illustration of the registry-shifted parallel N-strand β-sheet model, shown in more detail in Supplementary Figure S3. Black, white, and gray shading indicates that the sidechain is above the plane of the β-sheet, below the plane, or not expected to adopt a β-strand conformation, respectively. Selected residues are circled to highlight anticipated multisite occupancy and coexistence of residues with and without β-strand secondary structure. D) Overlays of 2D NMR contours from lyophilized samples (8 and 10; black) and centrifuged samples (A and B; red), showing spectra evidence of multisite occupancy. Solid and dashed rectangles indicate signal regions corresponding to β-strand and non-β strand secondary structures. E) Illustration of the planar antiparallel C-strand β-sheet model shown in more detail in Supplementary Figure S3, with black and white shading similar to panel C.

The “L”, “C”, and “2” symbols in Figure 1A correspond to the sample preparation methods used to detect inter-residue contacts via 2D ^13^C-^13^C NMR. The “0” symbols indicate pairs of residues for which inter-residue proximities were not detected by 2D NMR despite both residues being isotopically labeled in the same sample, as illustrated by the dashed circles in the 2D NMR spectra. We studied samples with new isotopic labels prepared using our established sample preparation method (36), and samples prepared with a modified method. The essential steps of the established protocol are: 1) isolation of monomeric Aβ42 by size exclusion chromatography (SEC); 2) induction of small oligomers called 2-4mers in the detergent SDS; 3) removal of SDS by dialysis, which induces further aggregation of Aβ42 to a 150 kDa size; 4) isolation of the 150 kDa oligomer by SEC; and 5) lyophilization and packing of the 150 kDa oligomer into solid-state NMR rotors. We refer to samples prepared with this protocol as “lyophilized” samples (see Supplementary Table S1), and contacts detected with lyophilized samples are indicated by “L” symbols in Figure 1A. Our modified sample preparation avoided lyophilization, instead employing ultracentrifugation (24 hours at 280,000g) to concentrate samples into NMR rotors. We refer to samples concentrated by centrifugation as “centrifuged” samples (see Supplementary Table S1). The “2” symbols indicate inter-residue proximities detected in lyophilized samples and subsequently confirmed in centrifuged samples. The “C” symbols indicate inter-residue proximities detected in centrifuged samples but not detected previously in lyophilized samples, likely because of the adverse effects of lyophilization-induced broadening on NMR sensitivity. While lyophilized samples can offer higher NMR intensities (more peptide in the NMR rotor), centrifuged samples exhibited sharper NMR lines, particularly when isotopic labels were on residues away from the centers of the β-sheets.

To our knowledge, there is no previously proposed structure or structural model that is consistent with our constraints for the 150 kDa oligomer. In particular, molecular configurations described in Supplementary Figure S1 predict exclusively in-register parallel (Figures S1A and S1B) or antiparallel (Figures S1C and S1D) β-sheets, and we show that parallel and antiparallel β-sheet coexist in the 150 kDa oligomer. In-register parallel β-sheets common to amyloid fibrils would anticipate no contacts beyond the gray-colored region of the contact chart without stacking of β-sheets via steric zippers, and thus cannot explain the pattern of contacts between residues *i* and *i* ± 3 or *i* ± 4 observed for the N-strand. For the C-strand, an antiparallel arrangement into β-sheets leads to a pattern of contacts along a direction perpendicular to the diagonal. This pattern is partially explained in Figures S1C and S1D.

We designed isotopic labeling schemes for the centrifuged samples to fulfill two purposes. First, we intended to test whether the oligomer structure within centrifuged samples matched that of lyophilized samples. Second, we sought to detect anticipated but previously undetected structural features. The 2D NMR results on centrifuged samples support the interpretation that centrifugation preserves the unique 150 kDa oligomer structure and reveals anticipated but previously undetected structural features. The “2” symbols in Figure 1A indicate contacts observed in lyophilized samples and verified in centrifuged samples. We chose isotopic labels to verify these contacts because they are unique to the 150 kDa oligomer structure (see Supplementary Figure S1). In addition we probed anticipated but previously undetected structural features, including multisite occupancy (coexistence of multiple molecular conformations) and inter-residue contacts foreseen by our modeling effort discussed below. One reason to expect multisite occupancy is the alternating registry-shifted alignment of neighboring N-strands, illustrated in Figure 1C. Using F19 as an example, Figure 1C shows that half of the F19 residues are proximal to K16 residues on adjacent β-strands and half are proximal to E22 residues on adjacent β-strands. The N-strand β-sheet structure further predicts 50% non-β-strand conformations for residues near the termini of the N-strands (e.g., V12 and D23). Since chemical shifts are most sensitive to local structures, occupancy of multiple conformations that do not undergo exchange would yield multiple NMR peaks for each isotopically labeled site. For previous results of lyophilized samples, we considered our spectral resolution too low to observe this effect. Figure 1D illustrates that, for centrifuged samples, NMR spectra exhibit the resolution of two Cα/Cβ crosspeaks for V12, F19, and D23. In this figure, dashed and solid boxes are used to indicate signals with ^13^C chemical shifts expected for residues with β-strand and non-β-strand secondary structures, respectively, supporting the notion that there are two inequivalent sites for V12, F19, and D23. Furthermore, as anticipated (Figure 1C), both F19 sites are within β-strands, whereas V12 and D23 each exhibit signals consistent with approximately 50% non-β-strand conformations. Additional support for multisite occupancy within the 150 kDa oligomer is found in the pattern of 2D NMR contacts between N-strand residues and C-strand residues (Figure 1A): for example, the contacts between F19 and I32 and between F19 and V36 seem to suggest that F19 is in two places at once, since I32 and V36 are at the edge and center, respectively, of the antiparallel β-sheet formed by the C-strand (see Figure 1E).

### Constraints on oligomer size, shape, and anisotropy from electron microscopy

Figure 2 shows EM imaging of the 150 kDa oligomer. Figure 2A is a negative-stain transmission EM image of 150 kDa oligomers isolated by SEC (see Supplementary Figure S4). Although these oligomers appear spherical in Figure 2A, they exhibit a tendency to assemble into “strings” (see Figure 2B). We estimate that strings represent only a minor part of the oligomer population: we observe them mostly when we aliquot from the high molecular weight shoulder of the 150 kDa SEC peak (see Supplementary Figure S4 and Table S3). The observation of strings suggests that oligomers form anisotropic interactions since spherically symmetric particles would not be expected to assemble in a preferential direction. As the oligomer is composed of β-sheets, we suggest that the anisotropy indicates preferred assembly along the β-sheet axis, possibly involving backbone hydrogen bonds. The strings appear to be self-limiting in their lengths, reaching up to ∼50 nm. Strings do not exhibit any evidence of helical symmetry. A similar size distribution of oligomer particles was observed in Aβ globulomers, which are also produced by adding SDS into solutions of Aβ monomers (39).

**Figure 2:**
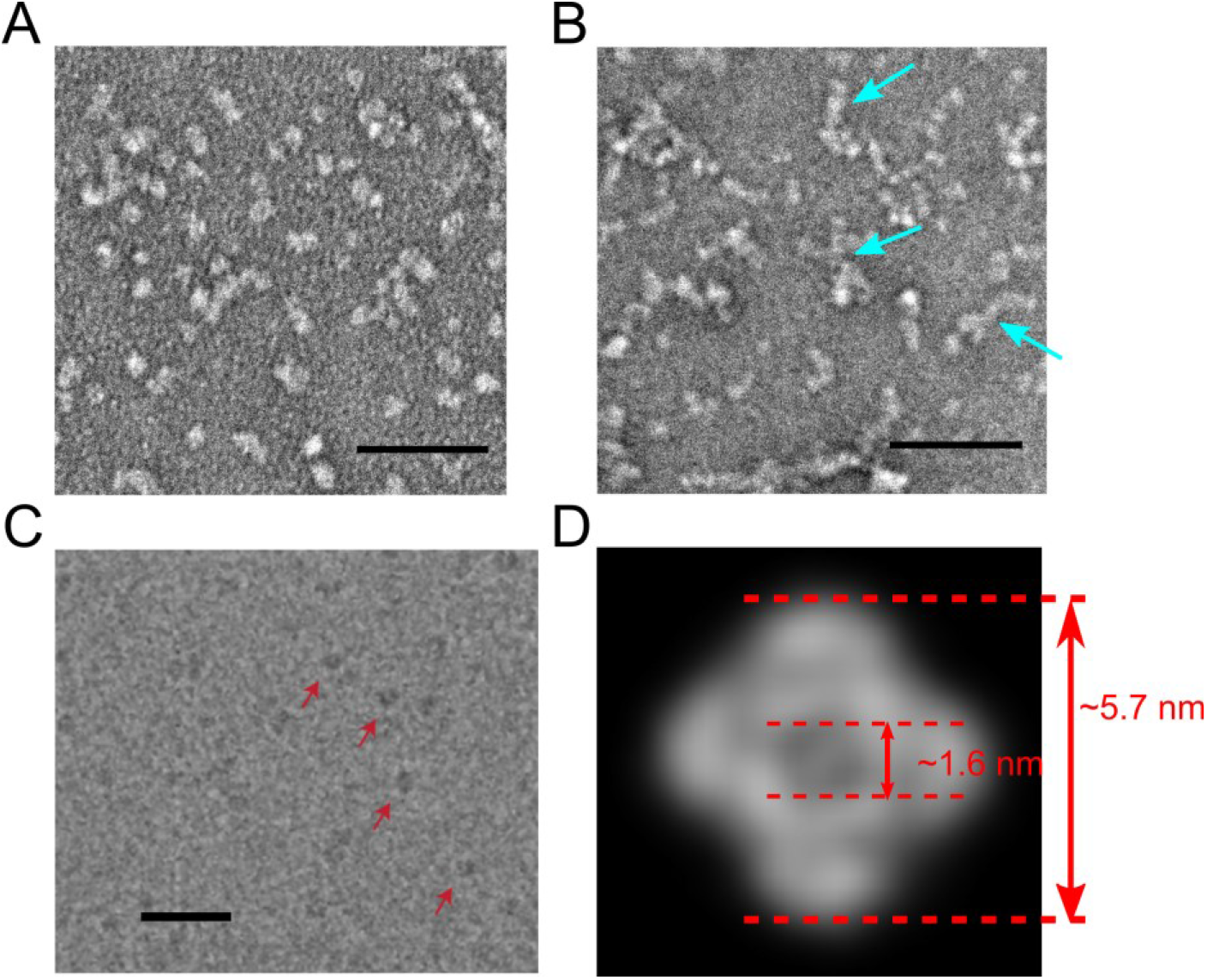
A) Negative-stain TEM image of 150 kDa oligomers. B) An image taken from the high-molecular-weight shoulder of the SEC peak, showing “strings” described in the text and indicated by arrows. C) Cryo-EM image of 150 kDa oligomers in ice, indicated by arrows. D) 2D cryo-EM reconstruction calculated from images like those in panel (C). Scale bars in (A-C) correspond to 50 nm.

For cryo-EM imaging, we froze aliquots from the center of the 150 oligomer SEC peak on C-Flat holey carbon grids and imaged with our Titan Krios equipped with a DE64 detector in electron counting mode. Embedded globular particles were clearly visible in the ice (Figure 2C), and a pore was visible in many of them. We performed cryo-EM single particle analysis on this species. We collected a dataset with Leginon to assess the quality of the sample, the distribution of particles in the ice, and the particle alignment. Contrast transfer function (CTF) estimates for selected globular particles were made, and 2D class averages were produced using cryoSPARC (Figure 2D). This analysis revealed a class average with four-fold symmetry and a clear pore in the center.

### Molecular structural modeling to harmonize NMR and EM constraints

The 4-fold symmetric 2D cryo-EM reconstruction (Figure 2D) inspired the following proposed β-strand arrangement, illustrated in Figure 3. This model is composed of 32 protomers (molecular weight 144 kDa) stacked into a hollow square cylinder shape (Figure 3A). Figure 3B illustrates the β-strand organization in each side of the square cylinder when it is cut open and unfurled (see dashed vertical line and scissor icon in Figure 3A). In this diagram, β-strands are drawn with green, magenta, blue, and red borders to mark four structurally equivalent “steric zippers,” each forming a side of the square cylinder (2 β-sheets and 16 β-strands per steric zipper). The β-strands are filled with gray or black coloring to indicate whether they are N-strands and C-strands. Half of the Aβ42 protomers have “U-shaped” conformations, such that both β-strands are in the same steric zipper. The other half of the protomers have “L-shaped” conformations (not apparent in the unfurled depiction). Each L-shaped protomer straddles between two neighboring steric zippers, enhancing oligomer stability. Figures 3C and 3D exhibit two cross-sections of our model, taken from adjacent layers perpendicular to the 4-fold symmetry axis, to highlight the difference between the U-shaped and L-shaped conformations, which alternate along the symmetry axis. Each diagram in Figure 3C and 3D is accompanied by a corresponding ribbon representation of the all-atom model. Curvature of C-strands is necessary to rationalize contacts with the N-strand and some of the contacts between C-strand residues.

**Figure 3:**
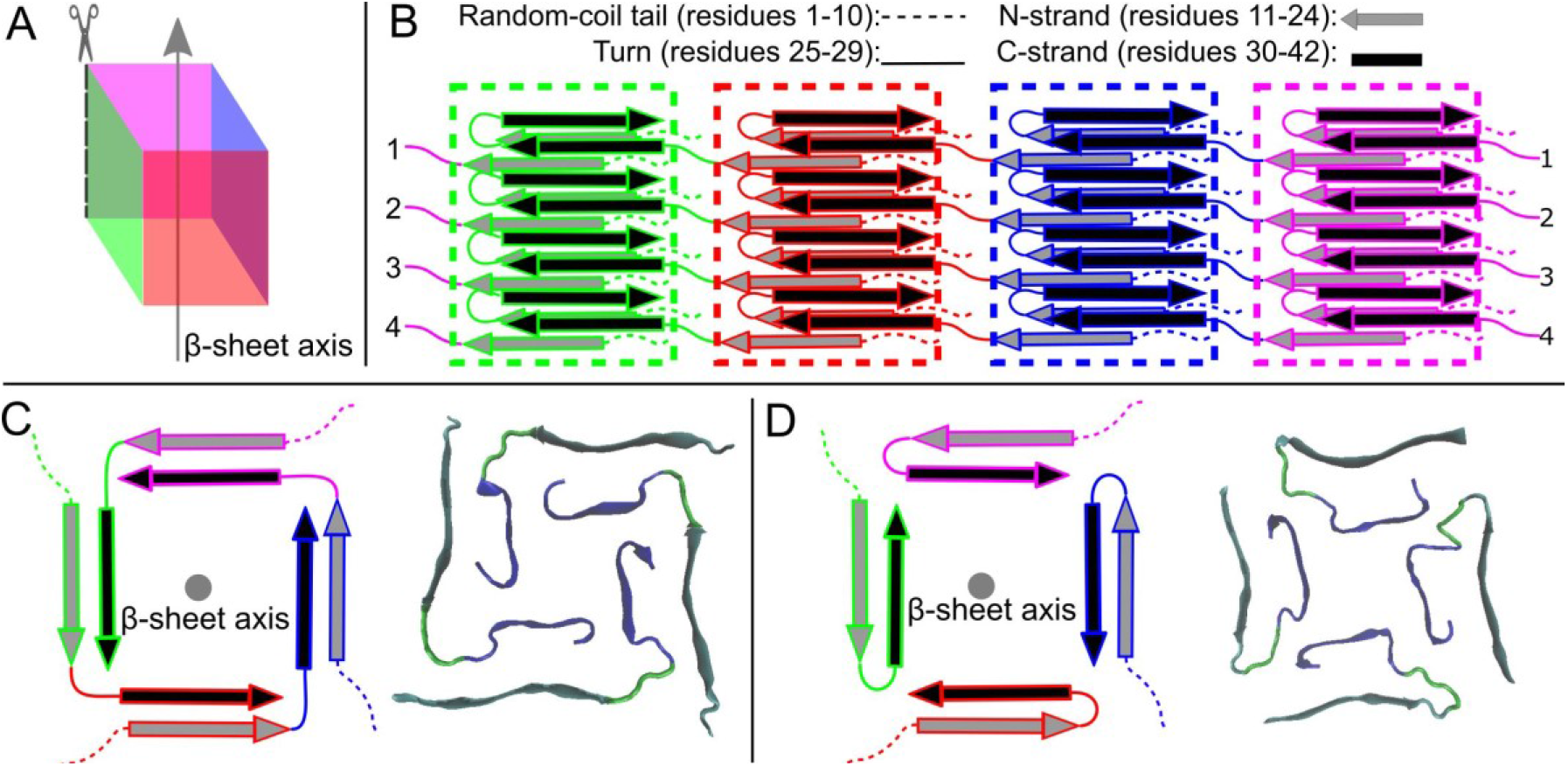
A) Proposed β-strand arrangement for the 150 kDa Oligomer. Four equivalent steric zippers are highlighted with green, red, blue, and magenta. Protomers are either in U-shaped or L-shaped conformations. Each U-shaped protomer contributes an N-strand and a C-strand to the same steric zipper, whereas L-shaped protomers domain swap such that its N-strand and C-strand are in distinct steric zippers. B) Schematic representation showing 3D organization of the steric zippers illustrated in panel A, drawn as if shape in panel A were cut along the dashed line and flattened. C and D) Depictions of adjacent layers sections perpendicular to the β-sheet axis. On the left of each panel is a schematic representation of β-strand orientations. On the right of each panel is a backbone presentation taken from our all-atom model. Layers are composed of protomers in U-shaped (panel C) or U-shaped conformations (panel D).

Figure 4 shows our final model and results of molecular dynamics simulations of the oligomer embedded in a membrane. Figure 4A exhibits the peptide backbones represented as ribbons. Figure 4B is the same inter-residue contact chart as in Figure 1A, but with a color scale corresponding to the model from Figure 4A. Based on previous measurements on isotopically diluted samples (34, 36), we estimate that the colored squares in Figure 4B correspond to pairs of residues for which proximity would be detectable by 2D ^13^C-^13^C NMR; however, this estimate is imprecise and likely to depend on ^13^C NMR line widths for the specific labeled sites. To improve agreement between the predicted and measured inter-residue contacts, we introduced two types of structural heterogeneity into the model. Inspired by the report of Ciudad et. al. on a Aβ42 tetramer (32), the model in Figure 4A includes molecules in β-hairpin conformations that cap the β-sheets, as well as heterogeneity in the alignment of C-strands. Supplementary Figure S5 describes more detail about our suggested heterogeneity in the C-strand alignment, and shows that this heterogeneity is consistent with our previous measurements of ^13^C-^13^C dipolar couplings between backbone carbonyl sites of V36. As shown in Figure 4B, our model rationalizes the 29 pairs of non-adjacent residues for which we detected inter-residue proximities (“L”, “C”, and “2” symbols in colored squares), and 28 out of 33 non-adjacent pairs for which proximity was not detected despite both residues being ^13^C-labeled within the same sample (28 “0” symbols in white squares and 5 in colored squares). The “0” symbols are individually inconclusive and would not normally be considered structural constraints in NMR modeling efforts, since the threshold of 2D NMR sensitivity is not precisely known. However, we consider it significant that our model rationalizes most of these negative results.

**Figure 4:**
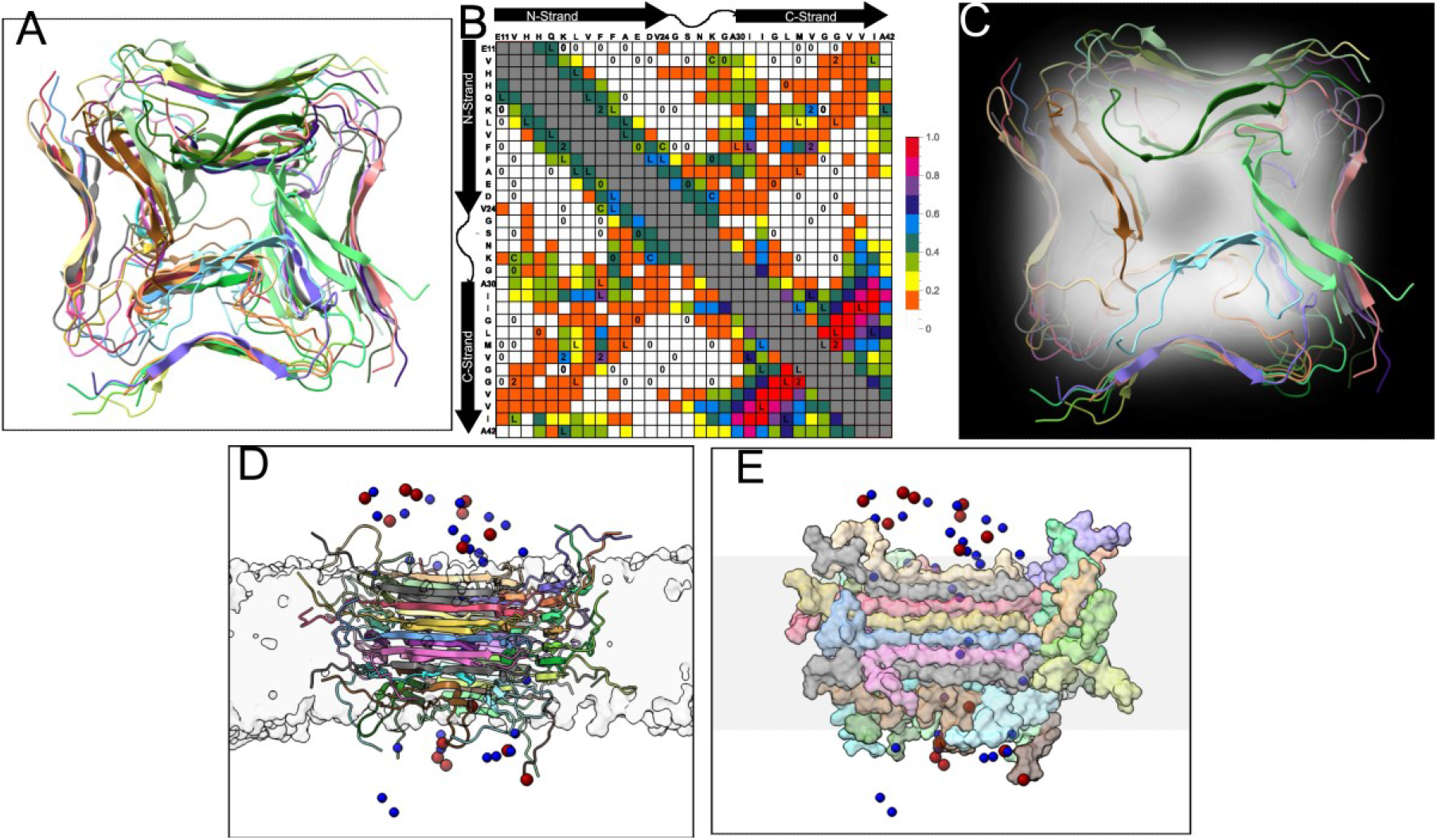
Results of model refinement and simulations in a membrane. To improve agreement between experimental and predicted inter-residue contacts, we introduced two types of disorder: capping of β-sheets with molecules in β-hairpin conformations, and disorder in C-strand alignment, as described in the text. We also performed unconstrained molecular dynamics simulations of the model in a membrane and simulations of ion transport. A) Refined model shown in cartoon representation, where chains are shown in different colors. Only residues 11-42 are shown. B) Contact chart based on the refined model. C) Overlay refined model structure with CryoEM density map. D) Oligomer embedded in a DOPC membrane. Proteins are in cartoon representation and colored according to chains. Ions are represented by sphere, with K+ and Cl- in blue and red, respectively. DOPC membrane is shown in light gray surface representation. E) Oligomer represented as surface, with chains colored in the same way as in (A); ions are as in (A); membrane is indicated by a gray band.

## Discussion

Based on our model, we suggest a reason why the 150 kDa oligomer does not undergo further aggregation into amyloid fibrils. We make the related suggestion that the 150 kDa oligomer is characterized by a fundamental tendency towards structural disorder that may distinguish oligomers from other types of aggregates. Although a level of structural order is indicated by harmony between our model and an array of experimental data, we suggest that the model may be too idealized and under-represent the diversity of structure within the 150 kDa oligomer samples. The model predicts perfect alternation between protomers in U-shaped and L-shaped conformations and is geometrically compatible with growth via molecular addition to fibril-like dimensions (not observed experimentally). However, while geometrically reasonable, such growth would be unlikely to occur. If Aβ42 monomer addition were to occur, newly added monomers would be unlikely to adopt the pattern of orientations and conformations necessary to propagate the structure. For similar reasons, we suggest that the 32-mer structure is unlikely to have been created by monomer addition to a smaller nucleus structure. Instead, we propose that this structure may be formed by the assembly of partially-ordered smaller oligomers. The tetramer structure reported by Ciudad et. al., for example, includes the organization of residues 29-42 into an antiparallel β-sheet resembling the C-strand within the 150 kDa oligomer (the precise C-strand alignment is partially identical). Perhaps this structure could undergo further assembly in which N-strands can organize into parallel β-sheets as tetramers interact. Consistently, we previously estimated, based on light scattering, SDS-PAGE, and circular dichroism measurements, that SDS initially induces the formation of 2-4mers, which, upon SDS removal, further aggregate into 150 kDa oligomers (33). TEM imaging suggests that 150 kDa oligomers can associate further into strings (see Figure 2B), which implies aggregation by oligomer assembly. However, growth into fibrils may be prevented by the supramolecular size of oligomers: while a monomer can undergo conformational changes as it docks onto a growing β-sheet, a 32-mer is unlikely to possess this type of plasticity. To our knowledge, our suggestion departs from previously proposed explanations of peptide oligomer limitation, which include barrel-like shapes (40, 41) and kinetic limitation via a dissociation rate that neutralizes the growth rate (42, 43).

In summary, our model may explain why oligomers could be stable without converting into fibrils. The model also suggests that it is possible for smaller oligomers to undergo further aggregation without forming fibrils. Finally, we predict that the 150 kDa oligomer could form an ion-conductive pore in a cell membrane.

## Acknowledgements

This work was supported by the National Institute on Aging of the National Institutes of Health and the National Institute on Minority Health and Health Disparities (award number RF1AG073434-01A1). The content is solely the responsibility of the authors and does not necessarily represent the official views of the National Institutes of Health.

## References

1. D. M. Walsh, D. J. Selkoe, Deciphering the molecular basis of memory failure in Alzheimer’s disease. Neuron 44, 181–193 (2004).

2. J. L. Tomic, A. Pensalfini, E. Head, C. G. Glabe, Soluble fibrillar oligomer levels are elevated in Alzheimer’s disease brain and correlate with cognitive dysfunction. Neurobiol. Dis. 35, 352–358 (2009).

3. M. L. Cohen et al., Rapidly progressive Alzheimer’s disease features distinct structures of amyloid-beta. Brain 138, 1009–1022 (2015).

4. S. T. Ferreira, M. V. Lourenco, M. M. Oliveira, F. G. De Felice, Soluble amyloid-beta oligomers as synaptotoxins leading to cognitive impairment in Alzheimer’s disease. Front. Cell. Neurosci. 9, 191 (2015).

5. E. N. Cline, M. A. Bicca, K. L. Viola, W. L. Klein, The Amyloid-beta Oligomer Hypothesis: Beginning of the Third Decade. J. Alzheimer’s Dis. 64, S567–S610 (2018).

6. S. Epelbaum et al., Acute amnestic encephalopathy in amyloid-beta oligomer-injected mice is due to their widespread diffusion in vivo. Neurobiol. Aging 36, 2043–2052 (2015).

7. C. A. H. Petersen et al., The amyloid beta-peptide is imported into mitochondria via the TOM import machinery and localized to mitochondrial cristae. Proc. Natl. Acad. Sci. U. S. A. 105, 13145–13150 (2008).

8. H. Du, S. S. Yan, Mitochondrial permeability transition pore in Alzheimer’s disease: cyclophilin D and amyloid beta. Biochim Biophys Acta 1802, 198–204 (2010).

9. E. J. Fernandez-Perez, C. Peters, L. G. Aguayo, Membrane Damage Induced by Amyloid Beta and a Potential Link with Neuroinflammation. Curr. Pharm. Des. 22, 1295–1304 (2016).

10. M. Mezler, S. Barghorn, H. Schoemaker, G. Gross, V. Nimmrich, A beta-amyloid oligomer directly modulates P/Q-type calcium currents in Xenopus oocytes. Br. J. Pharmacol. 165, 1572 (2012).

11. D. Hermann et al., Synthetic Abeta oligomers (Abeta(1-42) globulomer) modulate presynaptic calcium currents: prevention of Abeta-induced synaptic deficits by calcium channel blockers. Eur. J. Pharmacol. 702, 44–55 (2013).

12. E. Sturchler, A. Galichet, M. Weibel, E. Leclerc, C. W. Heizmann, Site-specific blockade of RAGE-Vd prevents amyloid-beta oligomer neurotoxicity. J. Neurosci. 28, 5149–5158 (2008).

13. M. Guglielmotto et al., Abeta1-42 monomers or oligomers have different effects on autophagy and apoptosis. Autophagy 10, 1827–1843 (2014).

14. T. Kawarabayashi et al., Dimeric Amyloid-β protein rapidly accumulates in lipid rafts followed by apolipoprotein E and phosphorylated tau accumulation in the Tg2576 mouse model of Alzheimer’s disease. The Journal of neuroscience : the official journal of the Society for Neuroscience 24, 3801 (2004).

15. W. K. Yu et al., Oligomerization of amyloid beta-protein occurs during the isolation of lipid rafts. J. Neurosci. Res. 80, 114 (2005).

16. A. K. Paravastu, R. D. Leapman, W. M. Yau, R. Tycko, Molecular structural basis for polymorphism in Alzheimer’s β-amyloid fibrils. Proc. Natl. Acad. Sci. U. S. A. 105, 18349 (2008).

17. A. K. Paravastu, A. T. Petkova, R. Tycko, Polymorphic fibril formation by residues 10-40 of the Alzheimer’s β-amyloid peptide. Biophys. J. 90, 4618 (2006).

18. W. Qiang, W. M. Yau, J. X. Lu, J. Collinge, R. Tycko, Structural variation in amyloid-beta fibrils from Alzheimer’s disease clinical subtypes. Nature 541, 217–221 (2017).

19. W. Qiang, W.-M. Yau, Y. Luo, M. P. Mattson, R. Tycko, Antiparallel β-sheet architecture in Iowa-mutant β-amyloid fibrils. Proc. Natl. Acad. Sci. USA 109, 4443–4448 (2012).

20. A. T. Petkova, W. M. Yau, R. Tycko, Experimental constraints on quaternary structure in Alzheimer’s β-amyloid fibrils. Biochemistry 45, 498 (2006).

21. M. T. Colvin et al., Atomic Resolution Structure of Monomorphic Abeta42 Amyloid Fibrils. J. Am. Chem. Soc. 138, 9663–9674 (2016).

22. L. Gremer et al., Fibril structure of amyloid-β (1–42) by cryo–electron microscopy. Science 358, 116–119 (2017).

23. U. Ghosh, K. R. Thurber, W. M. Yau, R. Tycko, Molecular structure of a prevalent amyloid-beta fibril polymorph from Alzheimer’s disease brain tissue. Proc. Natl. Acad. Sci. U. S. A. 118 (2021).

24. A. Abelein, J. Jarvet, A. Barth, A. Graslund, J. Danielsson, Ionic Strength Modulation of the Free Energy Landscape of Abeta40 Peptide Fibril Formation. J. Am. Chem. Soc. 138, 6893–6902 (2016).

25. E. Y. Chi et al., Amyloid-beta fibrillogenesis seeded by interface-induced peptide misfolding and self-assembly. Biophys. J. 98, 2299 (2010).

26. W. Qiang, W.-M. Yau, R. Tycko, Structural Evolution of Iowa Mutant β-Amyloid Fibrils from Polymorphic to Homogeneous States under Repeated Seeded Growth. J. Am. Chem. Soc. 133, 4018 (2011).

27. S. Chimon et al., Evidence of fibril-like β-sheet structures in a neurotoxic amyloid intermediate of Alzheimer’s β-amyloid. Nat. Struct. Mol. Biol. 14, 1157 (2007).

28. L. Yu et al., Structural characterization of a soluble amyloid beta-peptide oligomer. Biochemistry 48, 1870–1877 (2009).

29. M. Ahmed et al., Structural conversion of neurotoxic amyloid-β1-42 oligomers to fibrils. Nat. Struct. Mol. Biol. 17, 561 (2010).

30. C. Lendel et al., A hexameric peptide barrel as building block of amyloid-beta protofibrils. Angew Chem Int Ed Engl 53, 12756–12760 (2014).

31. D. Shea et al., alpha-Sheet secondary structure in amyloid beta-peptide drives aggregation and toxicity in Alzheimer’s disease. Proc. Natl. Acad. Sci. U. S. A. 116, 8895–8900 (2019).

32. S. Ciudad et al., Aβ(1-42) tetramer and octamer structures reveal edge conductivity pores as a mechanism for membrane damage. Nat. Commun. 11, 3014 (2020).

33. V. Rangachari et al., Amyloid-β(1-42) rapidly forms protofibrils and oligomers by distinct pathways in low concentrations of sodium dodecylsulfate. Biochemistry 46, 12451 (2007).

34. D. Huang et al., Antiparallel beta-Sheet Structure within the C-Terminal Region of 42-Residue Alzheimer’s Amyloid-beta Peptides When They Form 150-kDa Oligomers. J. Mol. Biol. 427, 2319–2328 (2015).

35. Y. Gao et al., Out-of-Register Parallel beta-Sheets and Antiparallel beta-Sheets Coexist in 150-kDa Oligomers Formed by Amyloid-beta(1-42). J. Mol. Biol. 432, 4388–4407 (2020).

36. W. M. Tay, D. Huang, T. L. Rosenberry, A. K. Paravastu, The Alzheimer’s Amyloid-β(1-42) Peptide Forms Off-Pathway Oligomers and Fibrils that are Distinguished Structurally by Intermolecular Organization. J. Mol. Biol. 425, 2494 (2013).

37. S. Matsumura et al., Two distinct amyloid β-protein (Aβ) assembly pathways leading to oligomers and fibrils identified by combined fluorescence correlation spectroscopy, morphology, and toxicity analyses. J Biol Chem. 286, 11555 (2011).

38. A. J. Dear et al., Kinetic diversity of amyloid oligomers. Proc. Natl. Acad. Sci. U. S. A. 117, 12087–12094 (2020).

39. S. Barghorn et al., Globular amyloid β-peptide(1-42) oligomer - a homogenous and stable neuropathological protein in Alzheimer’s disease. J. Neurochem. 95, 834 (2005).

40. M. Serra-Batiste et al., Abeta42 assembles into specific beta-barrel pore-forming oligomers in membrane-mimicking environments. Proc. Natl. Acad. Sci. U. S. A. 113, 10866–10871 (2016).

41. L. Connelly et al., Atomic force microscopy and MD simulations reveal pore-like structures of all-D-enantiomer of Alzheimer’s beta-amyloid peptide: relevance to the ion channel mechanism of AD pathology. J Phys Chem B 116, 1728–1735 (2012).

42. J. Yang et al., Direct Observation of Oligomerization by Single Molecule Fluorescence Reveals a Multistep Aggregation Mechanism for the Yeast Prion Protein Ure2. J. Am. Chem. Soc. 140, 2493–2503 (2018).

43. T. C. T. Michaels et al., Dynamics of oligomer populations formed during the aggregation of Alzheimer’s Aβ42 peptide. Nat. Chem. 12, 445–451 (2020).

