## Supplementary Information for "Structural Model for Self-Limiting β-strand Arrangement Within an Alzheimer’s Amyloid-β Oligomer"

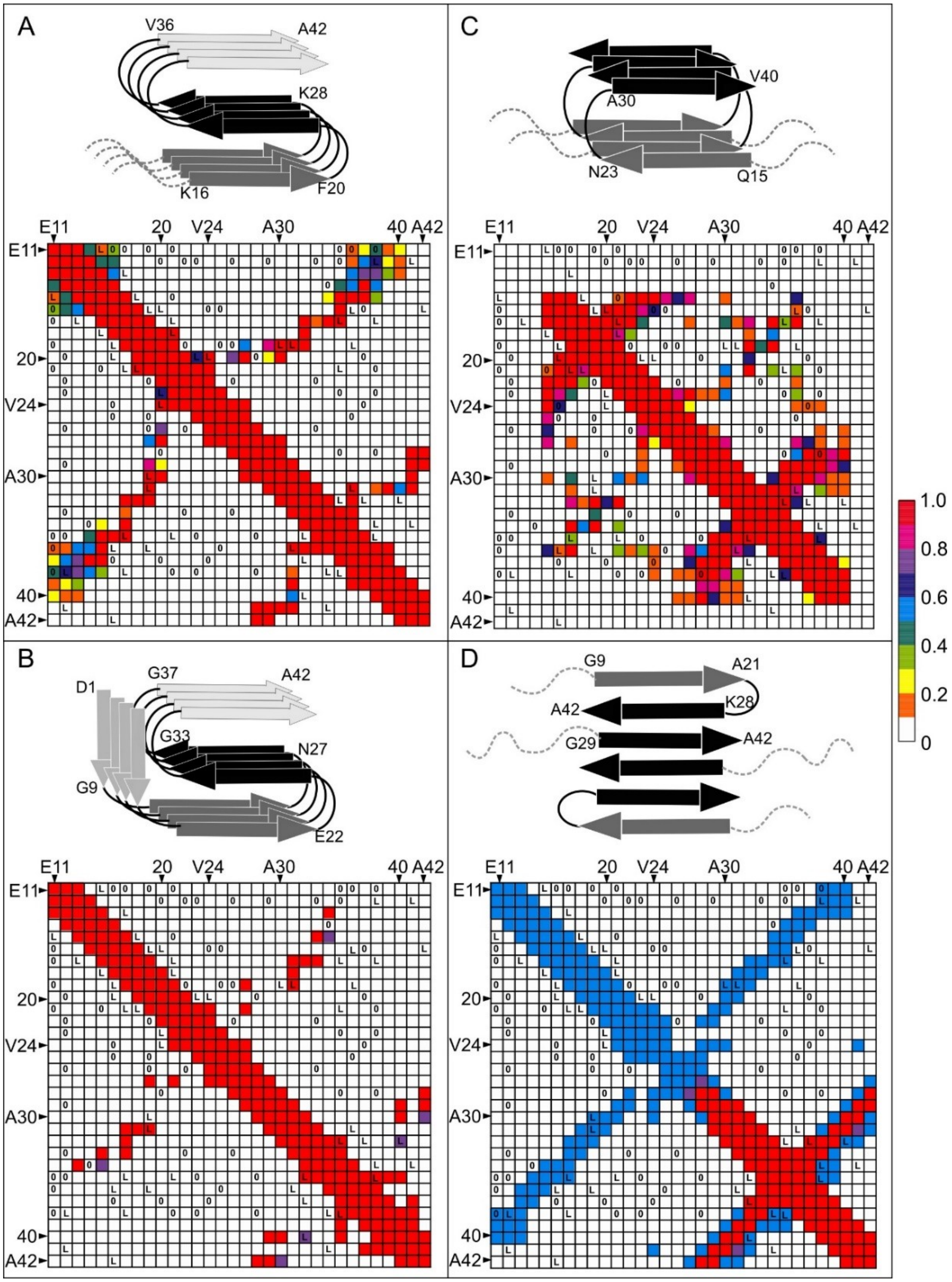


Figure S1: Contact charts for previous models, with symbols corresponding to our previously published inter-residue contacts for the 150 kDa oligomer. The "L" symbols indicate inter-residue contacts observed in 2D $^{13}C$-$^{13}C$ spectra on lyophilized samples. Each contact chart is accompanied by a schematic of the β-strand arrangement in the previously published structure or structural model. Panels A and B correspond to the Aβ(1-42) fibril structures reported by main text References 21 and 22, respectively. Panel C corresponds to the structural model for a fibril formed by the Iowa mutant of Aβ(1-40), reported by main text Reference 19. Panel D corresponds to the tetrameric Aβ(1-42) oligomer structure reported by main text Reference 32.

Table S1. List of 150 kDa oligomer samples analyzed by solid-state NMR. The samples differ in isotope-labeling scheme and sample preparation method. NMR spectra for lyophilized samples were presented in our previous work. Centrifuged samples were analyzed for the first time in the present work.

| Sample | Sample Preparation Method | Isotope Labeling (U-$^{13}C/^{15}N$, indicated residues) | Results Location or Citation |
|---|---|---|---|
| **A** | Centrifuged | V12, D23, K28, M35, G38 | Figures 1B and S2A |
| **B** | | K16, F19, G25, V36 | Figure S2B |
| **C** | | F19, E22, V24, S26 | Figure S2C |
| **1** | Lyophilized | K16, F20, V24, G37 | Main Text Reference 35 |
| **2** | | D7, G9, E11, L17, F19, A21 | |
| **3** | | E11, F19, I31, V36 | |
| **4** | | E11, L17, A21, M35, G38 | |
| **5** | | Q15, V18, A21 | |
| **6** | | S8, Y10, V12, L34, G38, I41 | |
| **7** | | V12, E22, S26, N27, G33 | |
| **8** | | V12, F20, D23, K28, G29 | |
| **9** | | E11, H13, Q15, L17 | |
| **10** | | E11, K16, F19, V36 | |
| **11** | | A2, E3, F4, G9, V39 | |
| **12** | | H14, K16, L34, A42 | |
| **13** | | I32, M35, G37, V40 | Main Text References 34 and 35 |
| **14** | | F19, V24, G25, A30, I31, L34, M35 | Main Text Reference 36 |

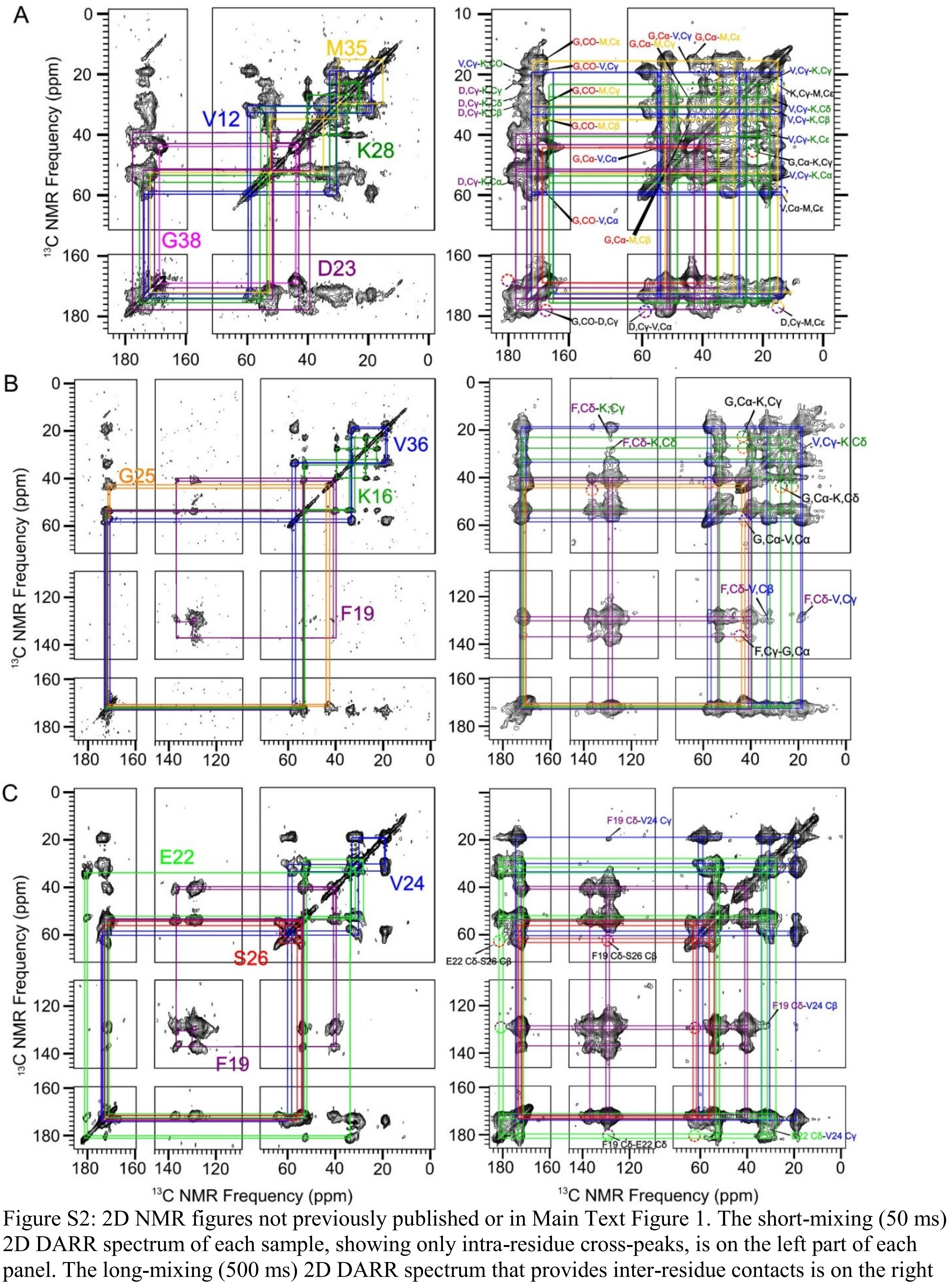


Figure S2: 2D NMR figures not previously published or in Main Text Figure 1. The short-mixing (50 ms) 2D DARR spectrum of each sample, showing only intra-residue cross-peaks, is on the left part of each panel. The long-mixing (500 ms) 2D DARR spectrum that provides inter-residue contacts is on the right part. A) sample A. B) sample B. C) sample C.

| Type | Residue | CO | $C^{\alpha}$ | $C^{\beta}$ | $C^{\gamma}$ | other C |
|---|---|---|---|---|---|---|
| ultra-centrifuge | V12 (β1) | 172.6/2.4 | 58.7/2.1 | 32.9/2.6 | 19.2/2.1 | |
| | | -2.0 | -1.8 | +1.7 | | |
| | V12 (c2) | 174.0/2.4 | 59.7/2.5 | 30.6/2.0 | 19.1/1.8 | |
| | | -0.6 | -0.8 | -0.6 | | |
| lyophilize | V12 | 174.1/3.2 | 58.8/4.0 | 32.7/2.9 | 19.3/3.3 | |
| | | -0.5 | -1.7 | +1.5 | | |
| ultra-centrifuge | K16 (β1) | 172.6/1.8 | 53.2/1.7 | 34.5/2.9 | 23.1/1.9 | $C^{\delta}$: 27.8/1.6, $C^{\epsilon}$: 39.9/1.3 |
| | | -2.3 | -1.3 | +3.1 | | |
| | K16 (β2) | 172.1/1.8 | 53.5/1.8 | 32.3/1.5±0.3 | 23.1/1.9 | $C^{\delta}$: 27.8/1.6, $C^{\epsilon}$: 39.9/1.3 |
| | | -2.8 | -1.0 | +0.9 | | |
| lyophilize | K16 | 171.3/3.7 | 53.2/2.6 | 34.4/5.2 | 23.9/2.2 | $C^{\delta}$: 28.0/3.0, $C^{\epsilon}$: 40.1/2.2 |
| | | -3.6 | -1.3 | +3.0 | | |
| ultra-centrifuge | F19 (β1) | 172.8/1.6 | 53.4/2.3 | 40.0/2.2±0.2 | 136.9/3.5 | $C^{\delta}$: 130.3/1.6±0.2, $C^{\epsilon}$: 130.0/1.6±0.2, $C^{\zeta}$: 128.7/1.4±0.2 |
| | | -1.3 | -2.6 | +2.1 | | |
| | F19 (β2) | 171.5/2.0 | 53.9/1.8 | 41.2/1.6 | 136.9/3.5 | $C^{\delta}$: 130.3/1.6±0.2, $C^{\epsilon}$: 130.0/1.6±0.2, $C^{\zeta}$: 128.7/1.4±0.2 |
| | | -2.6 | -2.1 | +3.3 | | |
| lyophilize | F19 | 172.1/2.7 | 54.2/3.9 | 40.8/2.2 | 136.6/3.2 | $C^{\delta}$: 129.0/2.6 |
| | | -2.0 | -1.8 | +2.9 | | |
| ultra-centrifuge | D23 (β1) | 170.6/4.6 | 51.4/2.1 | 42.8/3.7 | 177.6/3.3±0.2 | |
| | | -4.0 | -1.1 | +3.4 | | |
| | D23 (c2) | 174.3/5.1 | 51.8/3.9 | 39.2/3.4 | 177.6/2.8±0.2 | |
| | | -0.3 | -0.7 | -0.2 | | |
| lyophilize | D23 | 173.6/4.2 | 51.6/3.2 | 40.7/8.2 | 177.8/4.4 | |
| | | -1.0 | -0.9 | +1.3 | | |
| ultra-centrifuge | G25 (c1) | 171.6/4.0±0.2 | 42.7/2.8 | | | |
| | | -1.6 | -0.7 | | | |
| | G25 (c2) | 170.8/1.2 | 44.1/1.1 | | | |
| | | -2.4 | +0.7 | | | |
| lyophilize | G25 | 171.0/4.7 | 43.4/3.7 | | | |
| | | -2.2 | 0.0 | | | |
| ultra-centrifuge | K28 (c1) | 175.3/1.2 | 55.6/1.8 | 30.7/1.8 | 23.1/2.8 | $C^{\delta}$: 27.1/1.9 $C^{\epsilon}$: 40.1/1.3 |
| | | +0.4 | +1.1 | -0.7 | | |
| | K28 (β2) | 174.0/2.9 | 53.3/2.7 | 32.7/3.5 | 22.9/2.4 | $C^{\delta}$: 27.1/1.9 $C^{\epsilon}$: 40.1/1.3 |
| | | -0.9 | -1.2 | +1.3 | | |

| | | | | | | |
|---|---|---|---|---|---|---|
| lyophilize | K28 | 174.7/6.6 | 55.3±0.2/5.9 | 34.0/5.1 | 23.3/4.6 | $C^{\delta}$: 27.9/3.2<br>$C^{\varepsilon}$: 40.7/2.6 |
| | | -0.2 | +0.8 | +2.6 | | |
| ultra-centrifuge | M35 (β1) | 171.8/2.3 | 52.4/2.3 | 34.8/3.6 | 30.1/2.8 | $C^{\varepsilon}$: 15.5/2.7 |
| | | -2.8 | -1.3 | +3.6 | | |
| lyophilize | M35 | 171.8/2.1 | 52.7/2.4 | 35.4/3.5 | 30.4/1.6 | $C^{\varepsilon}$: 15.6/2.3 |
| | | -2.8 | -1.0 | +4.2 | | |
| ultra-centrifuge | V36 (β1) | 172.4/2.4 | 58.4/1.7 | 33.8/1.9 | 19.4/2.4 | |
| | | -2.2 | -2.1 | +2.6 | | |
| | V36 (β2) | 170.4/1.2 | 57.0/1.9 | 33.6/2.1 | 18.8/3.8±0.2 | |
| | | -4.2 | -3.5 | +2.4 | | |
| lyophilize | V36 | 172.4/3.0 | 57.6/3.1 | 33.6/2.9 | 19.3/2.9 | |
| | | -2.2 | -2.9 | +2.4 | | |
| ultra-centrifuge | G38 (β1) | 168.7/2.1 | 43.8/2.3 | | | |
| | | -4.5 | +0.4 | | | |
| lyophilize | G38 | 168.5/1.8 | 44.0/2.2 | | | |
| | | -4.7 | +0.6 | | | |

Table S2: NMR chemical shifts (ppm)/linewidths (full width half maximum, ppm) for all $^{13}C$-labeled sites in the Aβ(1-42) 150kDa oligomer samples A, B, and C. The carbons in each amino are labeled as they are in the Biological Magnetic Resonance Data Bank. Estimated error is ±0.1 ppm for both the chemical shift and line width unless specified otherwise.

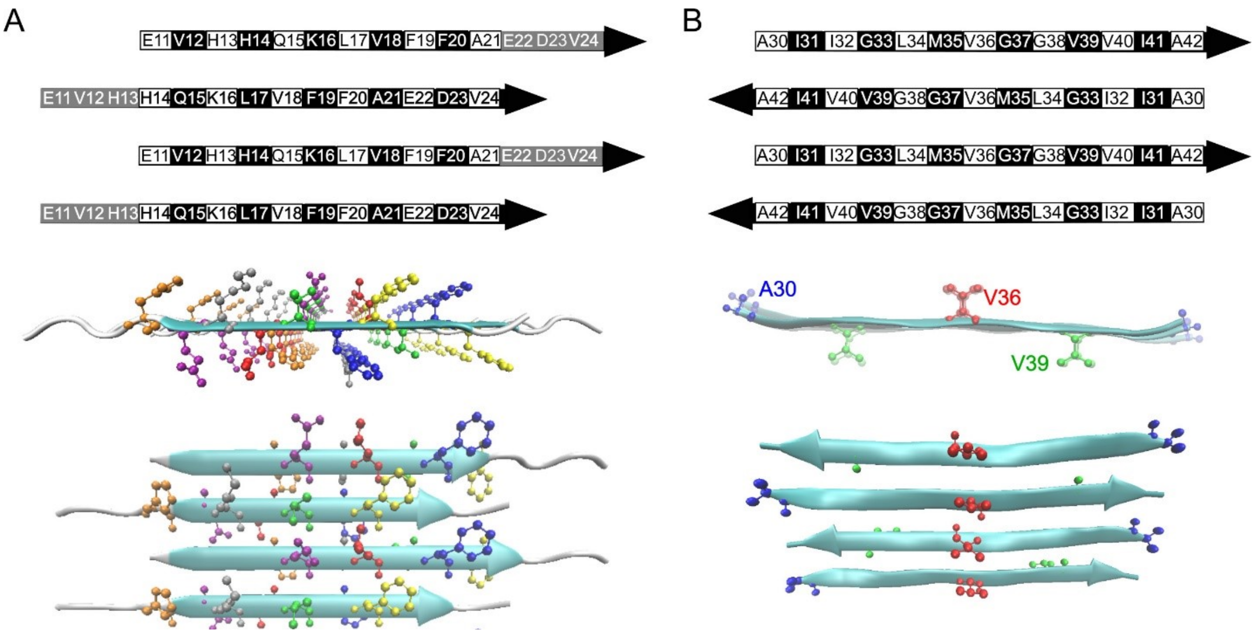


Figure S3: Depictions of N-strand and C-strand (Panels A and B, respectively) β-sheet models used to create the color scale in main text Figure 1A.

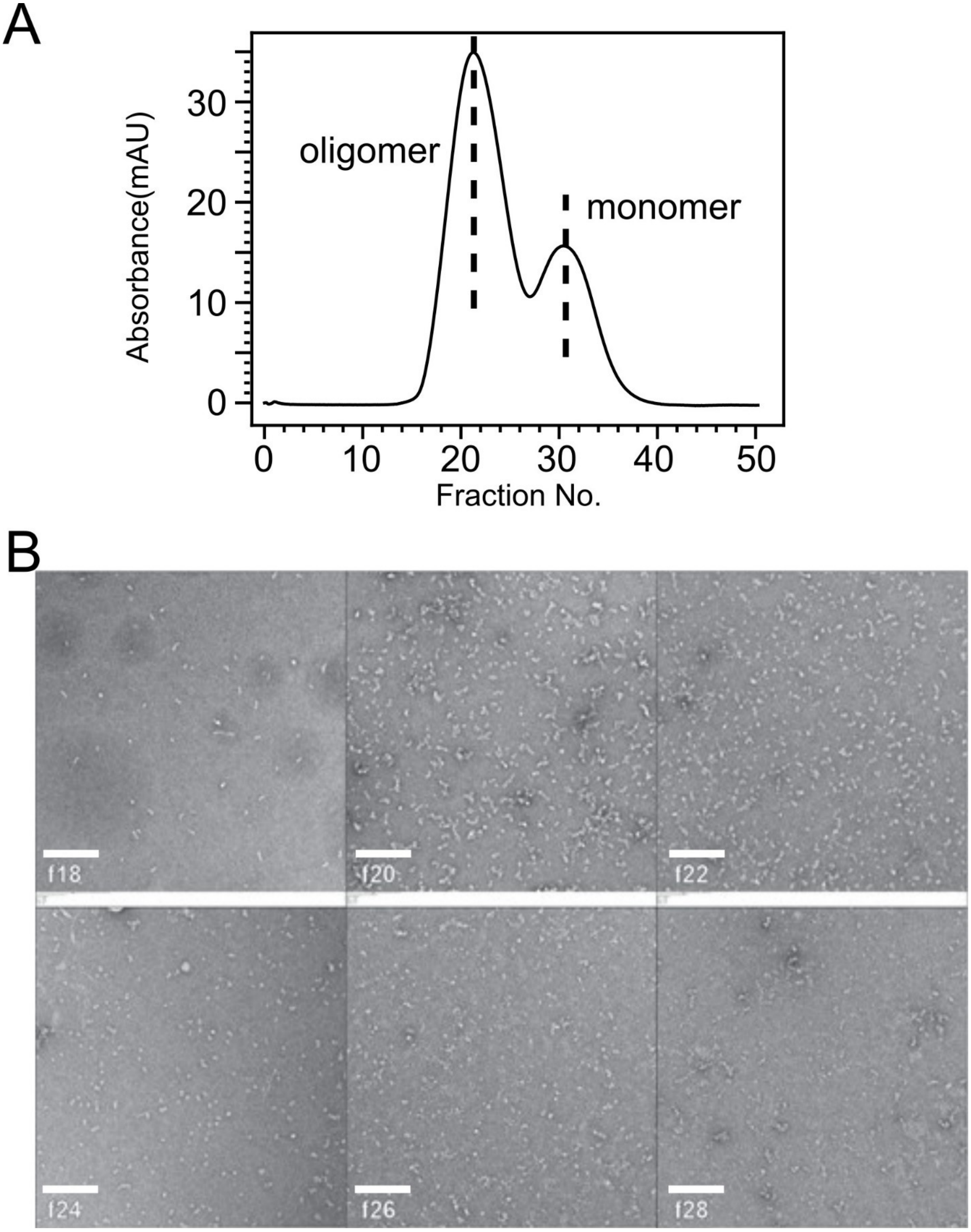


Figure S4: Panel A: SEC trace for 150 kDa oligomer solution. Panel B: TEM images taken from indicated SEC fractions.

| SEC Fraction: | f18 | f20 | f22 | f24 | f26 | f28 |
|---|---|---|---|---|---|---|
| Average (nm): | 19.9 | 25.3 | 14.9 | 12.1 | 13.1 | 12.1 |
| Std Dev (nm): | 27.1 | 10.2 | 5.7 | 4.5 | 4.7 | 3.8 |

Table S3. Estimation of the particle size distributions (based on diameters) of oligomers in different SEC fraction (Figure S4).

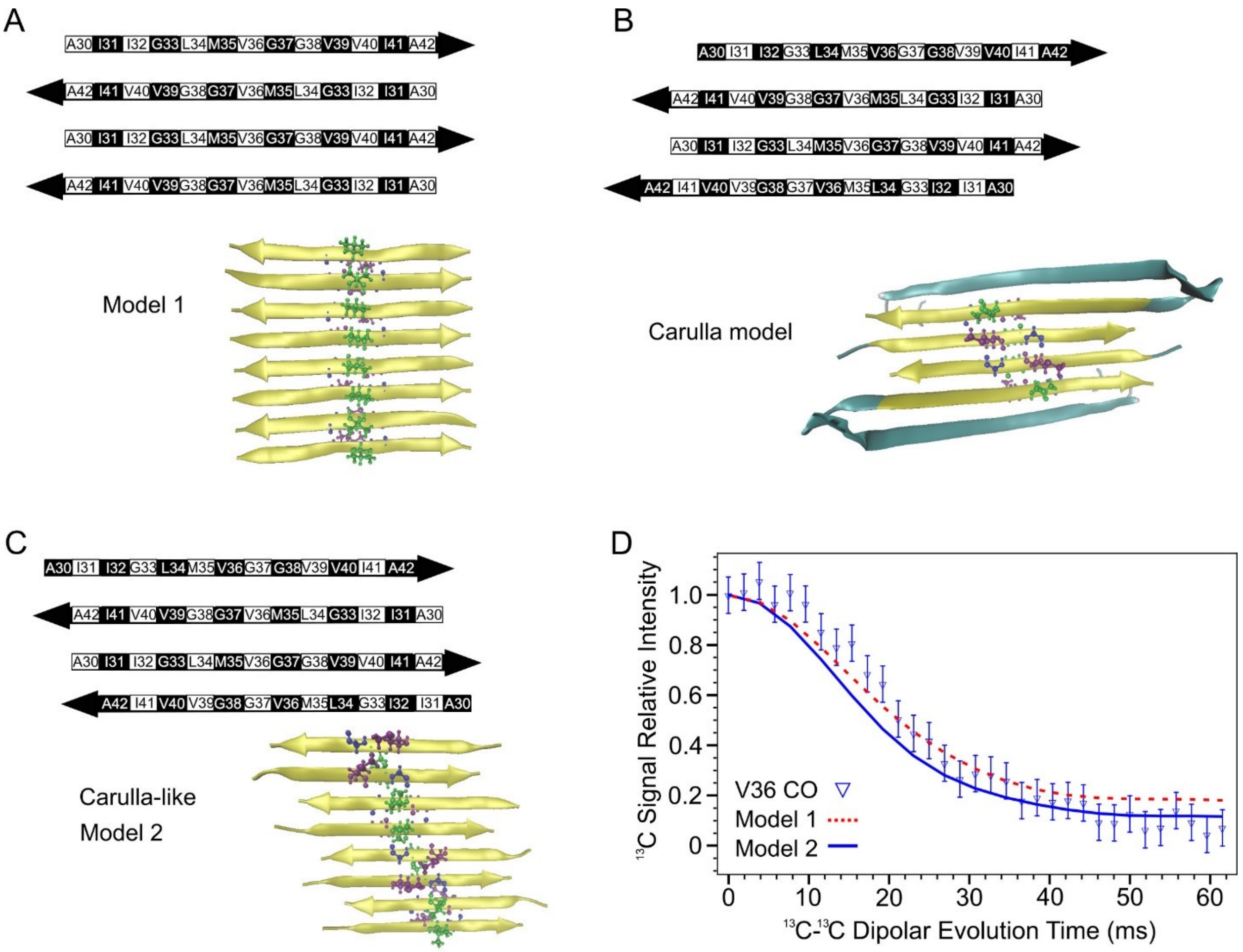


Figure S5: PITHIRDS-CT simulations showing that data can accommodate Carulla-like registry shifts in the C-strand alignment. A) A model of an antiparallel C-strand β-sheet centered at residue V36. B) A model of the C-strand β-sheet capped with β-hairpins, as predicted in the tetramer structure in main text Reference 32. C) An antiparallel C-strand β-sheet model in which the central residue alternates between residues G37 and V36. D) Simulated PITHIRDS-CT decays of $^{13}C$ NMR signal for a single $^{13}C$-label at the carbonyl carbon of V36 for the models in Panels A-C, compared to our previously published experimental data (main text Ref. 36).
